# Dynamic cell wall modifications in brassicas during clubroot disease

**DOI:** 10.1101/2020.03.02.972901

**Authors:** Julia Badstöber, Stefan Ciaghi, Sigrid Neuhauser

## Abstract

Biotic interactions of plants and microbial pathogens can cause drastic changes in cell wall composition in response to developmental reprogramming caused as consequence of an infection. Clubroot disease, caused by the biotrophic plant pathogen *Plasmodiophora brassicae* (Phytomyxea, Rhizaria), is the economically most important disease of Brassica crops worldwide. The disease is best known by the characteristic hypertrophied roots (root galls, clubroots). Amongst a series of physiological changes of the host tissue, the formation of the characteristic root galls leads to cell wall modification and reorganization. Cell wall chemistry and the hosts genetic repertoire are discussed to play a role in the resilience of plants against clubroot disease. Plant cells infected with *P. brassicae* are markedly enlarged, and look very differently from uninfected, healthy cells. Here we systematically review cell wall related processes that lead to the typical clubroot phenotype and provide novel insights how *P. brassicae* uses these modifications to benefit its own development. An infection with *P. brassicae* impacts on nearly all cell wall related processes, but all alterations are meaningful for successful growth and development of *P. brassicae.* Processes related to cell wall stability and rigidity (e.g. cellulose, pectin or lignin synthesis) are down-regulated, while cell wall degrading enzymes or processes that increase the flexibility of the host cell wall (e.g. expansin) are up-regulated. The here presented findings indicate that *P. brassicae* weakens the structural stability of its host cell while it increases its elasticity, which in consequence allows *P. brassicae* to grow bigger and ultimately to develop more resting spores. Consequently, the understanding of the modification of the host cell wall is important for the formation of the characteristic root galls but also to better understand clubroot disease.

## Introduction

Upon infection with the obligate biotrophic plant parasite *Plasmodiophora brassicae* (Phytomyxea, Rhizaria) marked morphological changes of the host cells occur. After infection the plant cells start to increase in size and start to alter their shape, which within a few weeks usually leads to gall formation in the roots (Kämper et al., 2006). These galls are the typical symptoms of clubroot disease (Figure 1). This morphological change goes inevitably hand in hand with marked changes in the transcriptomic profile of the host (Zhao et al., 2017;Irani et al., 2018;Olszak et al., 2019). Gall formation in the host is a common feature of phytomyxid parasites: 19 of the 42 phytomyxid species induce hypertrophies in their hosts (Neuhauser et al., 2011;Murúa et al., 2017). The induction of hypertrophies is not limited to plant hosts but can also be found in brown algae and oomycetes (Neuhauser et al., 2014). This formation of galls on the host is linked to the intracellular development of the phytomyxids: gradually larger areas of the host are infected, and multinucleate plasmodia develop and fill their host cells during growth. Phytomyxid growth and development goes hand in hand with developmental reprogramming of the host. Aim of this reprogramming is to generate a sink for nutrients and to create more space for the phytomyxid to grow and ultimately to produce as many resting spores as possible. Here we will discuss the developmental reprogramming of the host associated with the brassica cell wall in clubroot infected plants.

**Figure 1:**
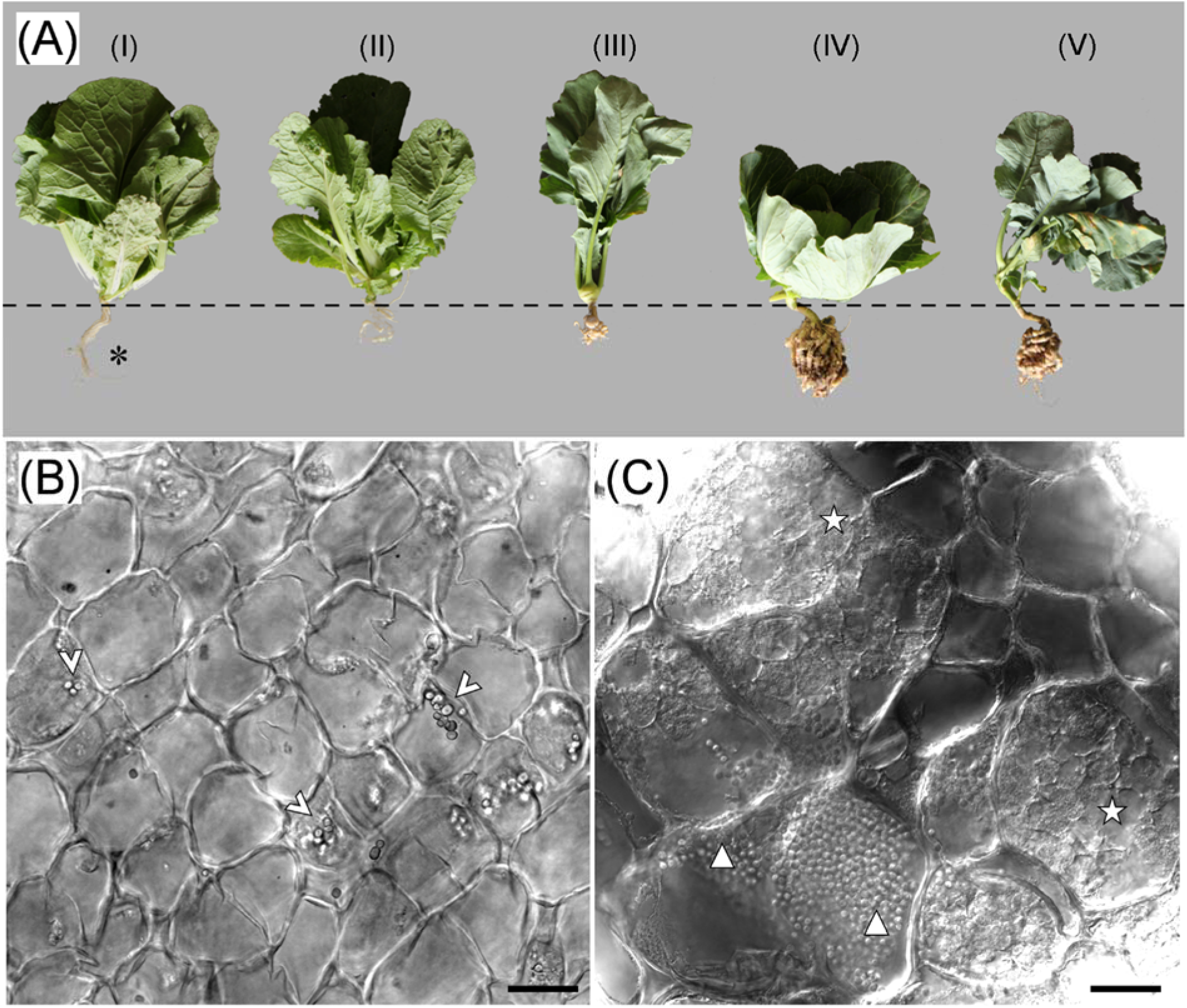
Clubroot disease in *Brassica* species. A: Typical clubroot symptoms caused by *Plasmodiophora brassicae*. Above ground development of infected plants is impaired while the roots show the typical hypertrophies. *Brassicae* spp. with and without (*)c clubroot symptoms. (I), (II) *Brassica rapa* subsp. *pekinensis* (Chinese cabbage); (III) *Brassica oleracea* var. *gongylodes* (kohlrabi); (IV) *Brassica oleracea* var. *capitata* (cabbage); (V) *Brassica oleracea* var. *italica* (broccoli). The dashed line indicates the soil surface. B: Section through a healthy *B. rapa* root. Starch granules are indicated with arrowheads. C: Section through a root gall of infected *B. rapa. Plasmodiophora brassicae* plasmodia are marked with stars and resting spores with triangles. Scale bar: 30 µm.

The plant cell wall mainly consists of polysaccharides and structural proteins (Figure 2A). The polysaccharides cellulose, hemicelluloses and pectin, are (in varying proportions) the major building blocks of plant cell walls. The currently accepted concept of cell wall structure describes the cell wall as (i) a reinforcing net like structure (“fishing net”) of cellulose microfibrils, which are held together by (ii) xyloglucans that bind to the net in clusters stabilizing it and (iii) a pectin gel consisting of stabilized, rigid parts (Ca^2+^) and very loose pectin gel structures (Cosgrove, 2016).

**Figure 2:**
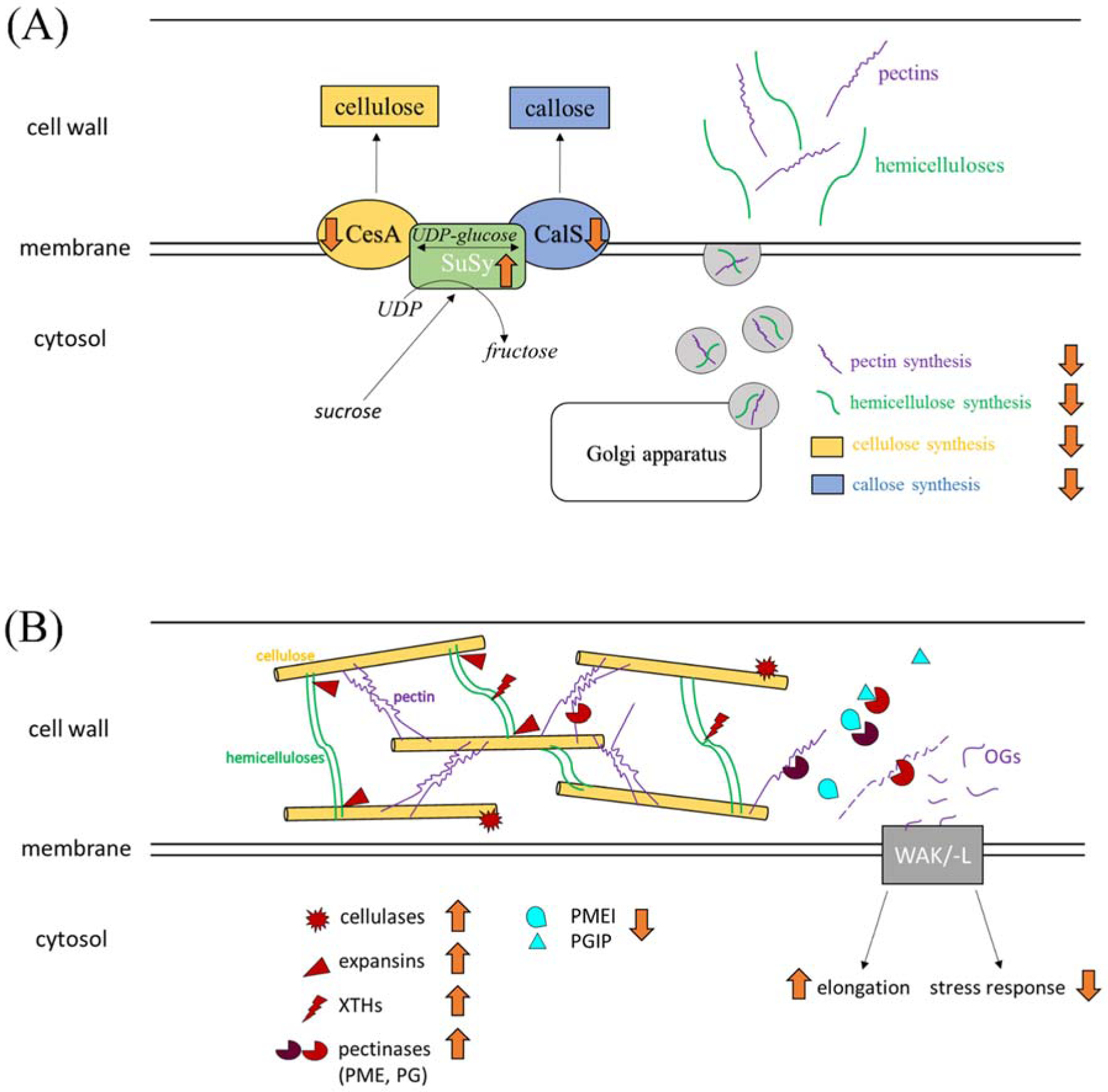
Synthesis of plant cell wall components (A) and modulation of cell wall components and related plant response (B). (A): Cellulose and callose synthesis are mediated through cellulose synthase (CeSA and callose synthase (CalS) in the plasma membrane. UDP-glucose serves as substrate for the synthesis. Sucrose synthase breaks down sucrose and generates UDP-glucose and fructose. Pectins and hemicelluloses are synthesized in the Golgi apparatus. Native pectins and hemicelluloses with sufficient length are transported by vesicles, secreted to the apoplast, and enzymatically incorporated into the cell wall. During infection with the biotrophic clubroot pathogen *Plasmodiophora brassicae*, genes involved in the synthesis of cellulose, hemicelluloses, pectins and callose are down-regulated. Genes coding for sucrose synthase are up-regulated in clubroots. Up- and down-regulated processes in clubroots is indicated with orange arrows. (B): Cell wall degrading and modulating enzymes and their target components. Cellulases modulate cellulose fibrils, XTHs cleave and re-anchor xyloglucan chains (hemicellulose), expansins disrupt linkages between hemicellulose and cellulose, and pectinases cleave pectins. Depolymerized pectins (oligoglacturonases, OGs) are bound and recognized by wall associated kinase/-like (WAK/L) and can trigger further plant response cascades. Pectinase-inhibitors (PMEIs, PGIPs) inhibit PME and PG activity. During infection with the biotrophic clubroot pathogen *Plasmodiophora brassicae*, genes like cellulases, pectinases (cell wall component degradation) and expansins and XTHs (cell expansion) are up-regulated in infected tissue. Enzyme inhibitors like PMEI or PGIP are down-regulated. Genes coding for WAKs (involved in cell elongation) are up-regulated and for WAKLs (stress response) are down-regulated in clubroots. Up- and down-regulated processes in clubroots is indicated with orange arrows.

Every plant cell has a primary cell wall, while secondary cell walls are formed in specialized areas of the plant. Primary cell walls are dynamic, growing structures. They are very dynamic in young cells and during cell division. The main function of primary cell walls is the protection, flexibility, basic structural support of the cytoplasm, but they also play a key role in the interaction between cells (Houston et al., 2016). Primary cell walls consist of carbohydrate-based polymers like cellulose, hemicellulose, pectins and glycoproteins (Cosgrove, 2005;Sarkar et al., 2009). Primary cell walls are relatively thin, flexible and strong in order to withstand growth related, high turgor pressure, to hold wall stress relaxation and to enable cell elongation (Hamant and Traas, 2010). Newly synthesized wall polymers become integrated by crosslinking, physical interactions or via enzymes. Once plant cells mature and once they have finished expansion and growth, the secondary cell wall biosynthesis is initiated resulting in a remodeled biosynthetic machinery (Nakano et al., 2015). In contrast to the primary cell wall, the secondary cell wall compounds are cellulose, hemicelluloses (mostly xylans) and lignin (Cosgrove, 2005;Sarkar et al., 2009). The secondary cell wall is thicker and is formed at a later stage of plant development. Secondary cell walls mainly provide rigidity and allow for upright growth (Houston et al., 2016). Compared to the primary cell wall, they provide tensile strength rather than extensibility (Speck and Burgert, 2011).

The water insoluble carbohydrate cellulose is the main building block of both, primary and secondary, plant cell walls. Cellulose forms the load-bearing structure of the cell wall, where its fibrous structure is responsible for structural integrity (Maleki et al., 2016). Cellulose is composed of β-1,4-linked glucan chains, which are organized by the cellulose synthase complex (CSC), located in the plasma membrane, and which crystallize to microfibrils (Desprez et al., 2007). The family of cellulose synthase genes (CesA, belonging to the glycosyltransferase family 2, GT2) are the best studied genes contributing to cell wall synthesis, probably because they are present in a multitude of eukaryotes and prokaryotes (Popper et al., 2011).

Cellulose is synthesised by the cellulose synthase complex which is located in the plasma membrane and made of several cellulose synthase (CesA) subunits (Jones et al., 2016) and a sucrose synthase (SuSy), which breaks down sucrose and generates UDP-Glucose (Figure 2A). Different isoforms of cellulose synthase proteins are responsible for the synthesis of cellulose during primary and secondary cell wall synthesis. CesA1, CesA3, and the CesA6-related proteins CesA2, CesA5, CesA9 are involved in primary cell wall cellulose synthesis, while CesA4, CesA7, and CesA8 proteins are responsible for secondary cell wall cellulose synthesis (Desprez et al., 2007;Li et al., 2016). CesAs which are involved in secondary cell wall biosynthesis, also confer resistance against bacterial and fungal pathogens. It is important to note that this resistance pattern is independent of salicylic acid, ethylene and jasmonic acid signaling (Hernandez-Blanco et al., 2007). It is suggested, that CesA1, CesA3 and CesA6 are present in the same plasma membrane protein complex responsible for cellulose biosynthesis, whereas CesA2, and CesA5 (related to CesA6) have partially redundant roles (Desprez et al., 2007). Mutation of any of these primary and secondary cell wall cellulose synthesis genes (CesA1, −3, −6 and CesA4, −7, −8) lead to modifications in composition and structure of both types of cell wall (Hernandez-Blanco et al., 2007;Kumar et al., 2018).

The cellulose synthase superfamily also contains nine cellulose synthase-like (Csl) families (CslA - CslJ). The Csl genes share similarities in sequence with the CesA genes and might be involved in the synthesis of the backbone of various polysaccharide polymers (Richmond and Somerville, 2000) like found e.g. in hemicelluloses of the plant cell wall (Lerouxel et al., 2006). Hemicelluloses include polymers like xyloglucans, mannans, glucomannans, and xylans and have β-1,4-linked backbones with an equatorial configuration (Scheller et al., 2010). Unlike cellulose, which is synthesized in the cell membrane, hemicelluloses and pectins are synthesized by glycosyltransferase (GTs) in the golgi apparatus (Suzuki et al., 2006) from where the native hemicellulose/pectins are transported to the membrane with vesicles. They are then secreted to the apoplast and enzymatically incorporated into the cell wall (Verbancic et al., 2018) (Figure 2B).

Pectins are the most heterogeneous and complex group of polysaccharides of the cell wall and mainly occur in the primary cell wall. Pectins found in the plant cell wall are homogalacturonan (HG), rhamnogalacturonan I (RGI) and II (RGII), and xylogalacturonan (XGA). Together with hemicelluloses they form the matrix in which cellulose microfibrils are embedded. Pectins play important roles for cell expansion, adhesion, strength, and porosity, but are also relevant for signaling between cells (Harholt et al., 2010;Ferrari et al., 2013). Together with other cell wall forming polymers, pectin is involved in providing physical strength to the plant. Plant pathogens can cause degradation of pectin by breaking it down into oligogalacturonides (OGs), which are involved in signaling for sensing pathogen caused wall degradation (Ferrari et al., 2013) (Figure 2B).

The deposition of lignin in the polysaccharide matrix during maturation of the secondary cell wall is typically the final step of secondary cell wall biosynthesis (Meents et al., 2018). It ensures two characteristic properties of the secondary cell wall: increased rigidity and impermeability. Lignin biosynthesis can be induced during cell development and maturation. But lignification of the plant cell wall can also be a response to biotic and abiotic stress, for example to protect other cell wall polysaccharides from degradation (Tronchet et al., 2010).

Plant cell walls are very dynamic, and responsive systems and can be modulated and remodeled by the plant itself (e.g. cell growth, elongation, reinforcement) or by pathogens (e.g. degrading enzymes (Underwood, 2012)). When *P. brassicae* infects the root cells of its host, those cells show a marked cell expansion, what differs significantly from cell growth in symptomless roots. Generally growth or an increasing size of plant cells is accompanied by the enlargement of the cell caused by an increase of turgor inside the plant which is generating cell wall stress (Cosgrove, 2016). This cell wall stress is relaxed by cell wall loosening processes (including xylanases, expansins etc) after a certain tipping point. These structural changes in the cell wall allow rapid expansion, which then again results in reduced turgor and the possibility for water to permeate the cell. This is followed by growth of the cell and by an increase of cell wall material. Cell wall stress is subsequently followed by reinforcement of the now “loose” cell wall through structural cell wall components. Enzymes like cellulases, Xyloglucan endotransglycosylases/hydrolases (XTHs) and expansins are mainly involved in wall expansion. They are supposed to disrupt noncovalent linkages between wall polysaccharides (Cosgrove, 2005) (Figure 2B).

Plant cell wall degrading enzymes (CWDEs) play an inevitable role for both, plants and pathogens. In plants they are involved in cell wall dynamics, e.g. in loosening, breakdown and the deposition of newly synthesized wall components (Marin-Rodriguez et al., 2002). They are also important for infection and establishment of plant pathogens (Kämper et al.). De-polymerization causes modification and alteration in wall rheology (Figure 2B). However, biotrophic pathogens like *P. brassicae* generally encode few CWDEs to avoid their recognition by the plant host (Kämper et al., 2006;Schwelm et al., 2015).

## Materials and Methods

Cell wall related genes were extracted from an existing *Brassica oleracea* var. *gongylodes* RNA-seq dataset (Ciaghi et al., 2019) were used (Accession: European Nucleotide Archive (ENA; https://www.ebi.ac.uk/ena) under the project PRJEB26435). These RNAseq data originated from white spindle galls and brownish spindle galls. Relevant pathways related to cell wall biosynthesis, rigidity, modulation, elongation, growth etc. were analyzed using all available information. Plant genes of interest (i.e. genes related with cellulose, hemicellulose, pectin, and lignin metabolism and cell expansion) were identified in this data using automatic gene annotations provided by MapMan (Thimm et al., 2004), KAAS (KEGG Automatic Annotation Server; Moriya et al., 2007), and eggNOG mapper (Huerta-Cepas et al., 2016). Transcripts fulfilling the criteria were curated manually. In general, only significantly differentially expressed genes (DEGs, adjusted p-value < 0.05) were analyzed but very rarely non-significant DEGs were included where it was useful for the biological interpretation and discussion of the data (noted in the text). This approach was chosen, because in a complex dataset like clubroots that contain a multitude of physiologically and developmental different cells smaller changes might still be relevant for a biological discussion of the plant pathogen interaction and for the formulation of a biological hypothesis. Log_2_-fold change values of significant DEGs (infected vs. control) and their corresponding FPKM (fragments per kilobase per million reads) values were used for this study, without any further threshold values.

MapMan, KAAS, eggNOG and TAIR Blast annotations were compared and genes grouped by those automated annotations. Putative isoforms were validated using alignments of the amino acid and protein sequence when contradictive expression values of DEGs were found. Isoforms were weighted based on their length, on sequence similarity and based on the general expression trend of all isoforms. Additionally, results published studies on clubroot disease were integrated to provide a comprehensive overview on cell wall related changes in clubroot infected plants.

## Results

### Synthesis of cell wall components

#### Cellulose synthase (CesA) and cellulose synthase-like (Csl) gene expression

Transcripts of the cellulose synthases CesA1-6 were identified in our data (Tab. S1). All of them were significantly down-regulated in clubroots compared with the control. CesA1-A6 transcripts were down-regulated during clubroot development.

Cellulose synthase-like A2 (CslA2) transcripts (Tab. S2) were significantly up-regulated in clubroots compared with the control, while CslA (Csl A3, 7, 11) transcripts were down-regulated, although this down-regulation was not statistically significant. CslA genes are involved in the beta-mannan biosynthesis, while CslC genes are supposed to be involved in xyloglucan synthesis. Of the CslC gene family transcripts belonging to CslC4, 5, 6, 8, 12 were identified in our dataset. Compared with the control, CslC4 and CslC6 was down-regulated in clubroots while CslC5 genes were up-regulated (Tab. S2). CslD genes are involved in the xylan and mannan biosynthesis and CslD3 were significantly down-regulated while CslD2, 5 were up-regulated. From the CslE family only CslE1 transcripts were significantly up-regulated clubroots.

To summarize: cellulose synthases (CesA family) were down-regulated in clubroots, whereas cellulose synthase-like genes (Csl) presented a more ambiguous picture, where some genes were up-regulated (CslA2, CslC5, CslD2, CslD5, CslE1) while others were down-regulated (CslA3, CslA7, CslA11, CslC4, CslC6, CslD3) in clubroot infected plants.

#### Pectin, lignin and callose synthesis genes

In clubroot infected tissue, transcripts linked to the synthesis of the most abundant pectins were down-regulated. Rhm1, Rhm2, and Quasimodo1 (Qua1) are coding for enzymes involved in the synthesis of rhamnose I (monosaccharides), and are involved in the homogalacturonan (polysaccharide) synthesis, two of the major components of pectin. All three genes were significantly down-regulated in clubroot tissue (Tab. S3).

Genes coding for an isoform of UDP-glucoronic acid decarboxylase (UXS), which produces UDP-xylose as a substrate for cell wall carbohydrates like pectin and hemicelluloses, were up-regulated (UXS1, UXS3, UXS5) and UXS2 was down-regulated in clubroot infected plants (Tab. S3).

In clubroot tissue, the genes involved in lignin biosynthesis were generally down-regulated in roots infected with *P. brassicae* (Tab. S4). The first three steps of the phenylpropanoid pathway, which initiates lignin biosynthesis are mediated through PAL (phenylalanine ammonia lyase), C4H (cinnamate 4-hydroxylase), and 4CL (4-coumarate:CoA ligase). Transcripts belonging to these gene families were down-regulated in clubroot infected tissue. Other lignin biosynthesis genes like COMT (caffeic acid O-methyltransferase), CCR (cinnamoyl-CoA reductase), and F5H (ferulate 5-hydroxylase) were also down-regulated.

Laccases and peroxidases are involved in the monolignol dehydrogenation step during lignin biosynthesis. With few exceptions genes coding for laccases and peroxidases involved in lignin synthesis were down-regulated in clubroot infected plants (LAC1, LAC3, LAC7, LAC11, LAC17). LAC6 and LAC13 were up-regulated.

Glucan-synthase-like genes (GSL), involved in callose biosynthesis, were generally down-regulated in clubroot tissue (GSL3, GSL7, GSL8, GSL10, GSL12) (Tab. S5). All these genes encode callose synthase (CalS). Genes coding for CalS1, the catalytic subunit 1 of callose synthase were up-regulated. UGT1 (UDP-glucose transferase 1), which is also involved in callose synthesis (transport of UDP-glucose to CalS), was down-regulated.

### 2. Degradation and modification of components

#### Cell wall degrading and modifying enzymes and inhibition of cell wall degradation

The most abundant transcripts of cell wall degrading enzymes found in clubroot infected plant tissue were pectinases and cellulases. Transcripts belonging to pectin degrading enzymes of the polygalacturonases (PGs), pectate lyases (PLs), pectin methylesterases (PMEs), and PAEs (pectinacetylesterases) were mostly up-regulated (some of them strongly) in infected roots (Tab. S7). Transcripts belonging to glycoside hydrolase family 28 proteins (GH28) were up-regulated in clubroots. All genes coding for PMEs (like PME3, PME 5, PME31), and most genes coding for PAEs (like PAE7, PAE 9-11) were up-regulated in infected plants, only PAE5 and PAE12 were down-regulated. Genes coding for callose degradation, or deposition were also up-regulated in clubroots (Tab. S5, S8).

Cellulases with hydrolase activity were generally up-regulated in clubroots, including glycosyl hydrolases family 9 (endoglucanases) like GH9B1, 6-8, 12, 13, GH9A1, and GH9C3. GH9B1, −13, GH9A1 and GH9C3 were down-regulated in root galls. Some genes of the glycosyl hydrolases family 17 were up-regulated in clubroots. Beta glucosidases were up- or down-regulated in clubroots (Tab. S8).

Inhibitors of cell wall degrading enzymes like polygalacturonases (PGs) and pectin methylesterases (PMEs), which dynamically modulate PMEs and PGs to inhibit pectin depolymerisation were down-regulated in clubroots (Tab. S9). Genes coding for PGIP2 (polygalacturonase inhibitor protein 2) were strongly down-regulated. PGIP1 were mainly down-regulated in infected roots. Genes coding for PMEIs (pectin methylesterase inhibitors) were also mainly down-regulated in clubroots, for example like PMEI6, −11 and PMEI13. Root hair specific 12 (RHS12) with pectin methylesterase inhibitor activity was also down-regulated in clubroots.

### 3. Cell wall loosening, elongation

#### *P. brassicae* infection causes cell wall loosening in plant roots

Genes involved in cell wall loosening and elongation processes like xyloglucan endotransglucosylases/hydrolases (XTHs) or expansins (EXP) and expansin-like genes (EXL) were mainly up-regulated (Tab. S10, S11). (XTH4, −9, −16, −24 and −32, EXP1, −6, −13, −15 and −20, EXPB2, and −3, EXLB1). In contrast genes like XTH 8, −10, and XTH21 and EXLA1, −2, and EXP 3 and EXP17 were down-regulated, in clubroots.

Wall associated kinases (WAKs) and wall associated kinase-like genes (WAKLs) are also involved in cell elongation, and all WAKs were up-regulated, whereas WAKLs were down-regulated in clubroots (Tab. S12).

Genes coding for SHAVEN3 (SHV3) and SHV3-like (SVL1) are also involved in cell expansion and alteration of wall rigidity. They were both up-regulated clubroots (Tab. S13).

## Discussion

### *Plasmodiophora brassicae* induces alterations of host cell wall rigidity

An infection with the phytomyxid *Plasmodiophora brassicae* results in molecular and physiological changes in the plant host: abnormal root growth results in gall formation. This is accompanied by altered metabolism, physiology and modifications of the host cell walls (Dekhuijzen and Overeem, 1971; Kavanagh and Williams, 1981; Ludwig-Müller et al., 1993; Devos et al., 2005; Ludwig-Müller et al., 2009; Malinowski et al., 2012). The growth of the host cells is resulting in surface increase, however, this is not automatically coupled to addition of new cell wall components (Cosgrove, 2014a). In clubroot infected cells the biosynthesis of wall components like cellulose, hemicelluloses, pectin (primary cell wall) and also lignin (secondary cell wall), which together form the load-bearing structures in the cell wall, is down-regulated (Tab. S1 – S4). These findings could be also shown in other experiments containing transcription analysis in clubroot infected *A. thaliana* (Irani et al., 2018). Genes involved in lignin biosynthesis are also down-regulated in former studies with infected *B. napus* (Cao et al., 2008; Ludwig-Müller et al., 2009). These results suggest an inhibition of cell wall component biosynthesis in clubroot infected tissue. The observed down-regulation of genes coding for biosynthesis of cellulose synthases (CesA), pectins (RHM, QUA) and hemicellulose (Csl genes) in infected plants supports this finding, which indicates that the cell walls of clubroot infected cells are less rigid than their counterparts in uninfected plants. Also transcripts of genes coding for cell wall degrading enzymes (CWDEs) like pectinases [polyglacturonases (PGs), pectate lyases (PLs) and pectin methylesterases (PMEs)] and cellulases with hydrolase activity were up-regulated (some strongly) in infected plants (Tab. S7, S8). These findings could also be validated in other studies of early infection of *B. oleracea* infected with *P. brassicae* (Wang et al., 2019).

In other words, *P. brassicae* is responsible for a decrease in root cell wall rigidity and stability in clubroots via down-regulation of the biosynthesis of primary and secondary cell wall components like cellulose, pectin, callose or lignin. On the other hand, enzymes which degrade these structural elements are up-regulated in clubroots. This loss in stability does fit the biological development of *P. brassicae*, because the multinucleate plasmodia gradually fill the significantly enlarged host cells. By reducing the rigidity of the cell wall, *P. brassicae* can grow to a larger size as the barrier posed by the cell wall gets more flexible. The processes seen at the transcriptomic level resemble the processes described during wall stress related growth as processes that control rigidity, but also extensibility is differentially regulated - with the main aim to withstand increasing turgor pressure without rupture of the cell (Cosgrove, 2016), while increased turgor-like pressure is built up by the growing parasite. That decreased rigidity of the cell wall has an important role for clubroot symptom development. This is supported by the finding that clubroot tolerant plants show transcriptomic signatures of an increased cell wall stability (Lahlali et al., 2017). The limited arsenal of CWDEs of *P. brassicae* has been notable when the genome of this parasite was analyzed (Schwelm et al., 2015;Rolfe et al., 2016). A lack of CWDEs is on the one hand a feature that is seen in many biotrophic plant parasites (Kämper et al., 2006). But on the other hand, the transcriptional changes observed in clubroot suggest that *P. brassicae* is able to compensate this by reprogramming the host metabolism.

In addition to a reduced rigidity of the plant cell wall strength, depolymerized pectin can serve as carbon source for pathogen development (Jia et al., 2009) and increases the accessibility of other plant cell wall components like cellulose and hemicelluloses to CWDEs, which can enable further and faster cell wall breakdown (Xue et al., 2018). This again would tie in very well with the biology of *P. brassicae*, which is obtaining nutrients from its host. So an increased availability of cell wall degradation products (sugars) can provide energy for *P. brassicae*. In order to protect the plant cells from excessive degradation of wall components through pectinases like PMEs or PGs, the host is able to produce enzyme-inhibiting enzymes, called PMEIs (PME-inhibitor) and PGIPs (PG-inhibiting protein). These enzymes modulate PMEs and PGs dynamically, and genes coding for these inhibitors were all down-regulated in clubroot tissue (Tab. S9). So as expected from a biotroph, *P. brassicae* induces changes in its host via inhibiting a cell wall related plant defense mechanism.

Plant genes involved in the metabolism of sugars that are building blocks of cell wall components were up-regulated in infected roots (Tab. S6). Transcripts of genes coding for sucrose synthase (SuSy), which catalyzes the reaction from sucrose to UDP-glucose and fructose are one example. UDP-glucose is needed as substrate for cellulose synthesis using CesA and for callose synthesis using CalS (Schneider et al., 2016) (Figure 2A). Genes coding for UDP-glucuronate decarboxylase (UXS), which produces UDP-xylose, a substrate for hemicelluloses and pectin, are up-regulated in clubroot transcriptomes (Tab. S3). Up-regulated sugar metabolism could also be shown in several other studies of clubroot infected *A. thaliana* (Siemens et al., 2006;Siemens et al., 2011;Walerowski et al., 2018). Summarizing these findings, it can be hypothesized that the increase in plant transcripts coding for CWDEs likely results in increased sugar availability via the degradation of cell wall components. On the other hand, the increase of transcripts belonging to sugar metabolic pathways likely results in an increased production of cell wall related sugars such as UDP-glucose or UDP-xylose. Overall it can be assumed, that these sugars might be taken up by *P. brassicae* as energy source, however, this hypothesis needs to be addressed separately.

### *Plasmodiophora brassicae* causes cell elongation and inhibits host defense response

To allow for cell expansion, an organized disruption of the existing cell wall homeostasis is followed by reinforcement via the formation of new linkages of wall components. Organized cell wall disruption can happen with the aid of CWDEs as described above, or via the enzymatic modification of wall rigidity and stability with expansins and xyloglucan endotransglycosylases/hydrolases (XTHs). Expansins and XTHs are directly involved and responsible for such modifications (Figure 2B). Transcripts of genes coding for expansins and for XTHs were significantly up-regulated in our data and previous transcriptome profiling works of clubroots (Siemens et al., 2006; Irani et al., 2018) (Tab. S10, S11). XTH activity was also detected in *B. rapa* by Devos et al. (2005). XTHs have two functions, either as xyloglucan endohydrolase (XEH), which cleaves xyloglucan chains, or as xyloglucan endotransglycosylase (XET), which is able to re-anchor two xyloglucan microfibrils, which have ruptured as result of turgor driven cell wall disruption (Rose et al., 2002). The majority of *B. oleracea* genes that were up-regulated in clubroots were annotated as XTHs possessing both functions (cleavage and re-anchoring). Notably, transcripts of genes that only feature the re-anchor activity (XET) were down-regulated. In other words, when *P. brassicae* plasmodia are growing in the infected host cell they might activate certain aspects of turgor driven cell growth in their host (up-regulation of expansins and XTHs with XEH and XET activity), but inhibit the plants ability to re-inforce the cell wall (down-regulation of XTHs with only XET activity). Other genes involved in cell expansion and alteration of wall rigidity with important roles in cell wall organization are genes coding for SHV3 (SHAVEN3) and SVL1 (SHV3-like). *Arabidopsis thaliana* SHV3 and SVL1 double mutants resulted in further defects like abnormal cell expansion, bloat guard cells as well as ectopic lignin deposits. Also pectin modifications and altered cellulose contents were observed (Hayashi et al., 2008). Related genes were up-regulated in clubroots (Tab. S7, S8, S13).

Wall associated kinases (WAKs) belong to the receptor-like kinases with a cytoplasmic protein kinase domain. They are located in the plasma membrane and serve as pectin receptors (cell wall side), for longer pectins of native cell walls and short pectic fragments of oligogalacturonic acid (OGs) (Kohorn, 2016) (Figure 2B). WAKs do not just bind pectins, they also recognize pectin fragments, which can be produced by pectin-degrading plant pathogens. This gene family is also involved in triggering stress response cascades in the host plant (Sivaguru et al., 2003; Kohorn and Kohorn, 2012). WAK genes are also discussed to be involved in cell elongation, because WAK mutants with lower levels of WAK proteins led to loss of cell expansion (Lally et al., 2001), shorter roots and a reduction in vacuolar invertase what might control expansion indirectly through turgor pressure (Kohorn et al., 2006; Kohorn et al., 2009; Kohorn and Kohorn, 2012). WAK-like genes (WAKL) share sequence similarities with WAKs, but have a direct role in disease resistance at least in maize (Hurni et al., 2015;Zuo et al., 2015). Evidence for this is provided, because dominant mutant allels of WAKLs in *A. thaliana* showed resistance to *Fusarium* (Diener and Ausubel, 2005). In clubroot infected plants, all genes coding for WAKs were up-regulated which fits very well to the above discussed increase in cell expansion. Overall, stress response is down-regulated in clubroots (Zhao et al., 2017), so an influence in stress response is not clear. All genes encoding WAKLs are down-regulated in clubroots, what might influence resistance negatively of host plants, because WAKLs are supposed to play a role in pathogen resistance (Verica and He, 2002) (Tab. S12).

Upon pathogen derived wall damage, the plant immune system can initiate a reinforcement of the cell wall as first line of defense, where a reinforced cell wall acts as physical barrier to infection. In order to slow down or to prevent pathogen invasion, callose or lignin containing cell wall thickenings are formed at infection sites (Stone, 1992). The most notable biochemical difference between clubroot-susceptible and resistant plants is an increased biosynthesis of lignin and phenolics, in the resistant plants (Lahlali et al., 2017). The key precursor of the phenylpropanoid and lignin pathways is L-phenylalanine (Barber and Mitchell, 1997; Shadle et al., 2003). PAL (L-phenylalanine ammonia-lyase) is the entry point enzyme to these pathways and the key regulator of the phenylpropanoid pathway, which can also be induced by pathogens (MauchMani and Slusarenko, 1996; Cozzolino, 2014). PAL and the lignin biosynthetic pathway appear to play an important role in resistance to clubroot conferring to the characterized clubroot resistance gene *Rcr1*. In clubroot resistant varieties of oilseed rape, PAL was highly up-regulated resulting in changes of the cell wall composition and lignification after contact with *P. brassicae* (Lahlali et al., 2017). PAL genes and the other important genes involved in lignin biosynthesis (coding for cinnamate 4-hydroxylase (C4H) and 4-coumarate:CoA ligase (4CL)) were down-regulated (Tab. S4), suggesting *P. brassicae* infection suppresses induced defense and resistance response of the *Brassica* host (Ciaghi et al., 2018).

Another possibility of cell wall reinforcement are callose-containing papillae (Voigt, 2014). Increased callose deposition is associated with the clubroot resistance gene *Rcr1* (Chu et al., 2014). Callose is synthesized in specialized cell walls like the cell plate (separates dividing cells) by the callose synthase, which is encoded by CalS genes belonging to the GSL (glucan syanthase-like) gene family. This family is divided in genes involved in fertility and cell division (GSL1, GSL2, GSL6, GSL8, GSL10) and in wall reinforcement (GSL5, GSL7, GSL12) (Verma and Hong, 2001). In our data, many transcripts could be annotated as GSL1, GLS8, GSL10 (fertility and division), and GSL7, GSL12 (reinforcement). All of them were down-regulated in our dataset (Tab. S5). GSL5, also known as powdery mildew resistant 4 (PMR4) involved in callose biosynthesis, is known to be re-located in the plasma membrane via vesicle-like bodies to papillae formation sites in *Arabidopsis thaliana* (Flors et al., 2008; Ellinger et al., 2013). GSL5 is (not significantly) down-regulated in our dataset, however, all available data suggest that an infection with *P. brassicae* leads to a decreased callose biosynthesis, deposition of callose synthase to pathogen site and for increased deposition of already synthesized callose, in order to reduce plant defense response. The genes PEN2 and PEN3 (Penetration2, 3) are also involved in callose deposition and glucosinolate activation and therefore necessary for plant resistance (Clay et al., 2009). Genes coding for PEN2 and PEN3 are up-regulated in our data, suggesting increased callose deposition and glucosinolate activation (Tab. S5). Glucosinolates are sulfur-containing glucosides and known as essential antimicrobial components in *Arabidopsis thaliana* (Hogge et al., 1988). Microorganisms are inhibited by breakdown products of glucosinolates derived through thioglucosidases (Kirkegaard et al., 1996; Manici et al., 1997; Tierens et al., 2001). In our data, genes coding for thioglucosidases, like TGG1 and TGG2 are extremely down-regulated, and thus hydrolysis into toxic, antimicrobial components (Tab. S14). This fact also suggests the pathogen *P. brassicae* inhibits plant defense response through suppression of the production of toxic antimicrobial substances.

## Conclusion

An infection of brassicas with the biotrophic plant pathogen *Plasmodiophora brassicae*, often leads to characteristic hypertrophied roots, the so called clubroots. Amongst other physiological changes clubroot infection leads to an altered biosynthesis of cell wall components. Genes coding for proteins involved in cellulose, hemicellulose, pectin, callose and lignin biosynthesis like CesAs (cellulose synthase), Csls (cellulose synthase -like), and CalS/GSLs (callose synthase) were down-regulated in infected plants. Also genes involved in pectin synthesis like Rhm and Qua1, or in the initial steps of lignin biosynthesis (PAL, C4H, 4CL) were down-regulated. Clubroot infection also leads to degradation and modification of existing wall components what results in decreased wall rigidity. Transcripts belonging to degrading/modifying enzymes, e.g. of pectins (PGs, PLs, PMEs, PAEs), cellulose through cellulases with hydrolase activity, or callose through degradation and deposition were up-regulated in clubroots. Gene expression for enzymes with degradation inhibiting activity were down-regulated in infected tissue. Clubroot infection also leads to massive host cell expansion, what we also see in transcriptomic data. Genes coding for XTHs and expansins, which allow cell wall loosening were up-regulated. Also for WAK/L genes, which are involved in cell elongation and defense response, gene expression differed. Elongation processes were up-regulated and defense response was down-regulated in clubroots. Genes involved in plant defense mechanisms leading to reinforcement of cell walls (e.g. callose synthesis, lignification) were down-regulated in clubroots, whereas deposition of callose was up-regulated. All these processes concur with observations during clubroot development, and the typical clubroot phenotype is a result of efficient strategies of *P. brassicae* to influence host biology. How and via which mechanisms this takes place remains to be answered with more targeted studies in the future.

## Acknowledgements

JB, SC and SN were funded by the Austrian Science Fund (FWF): grant Y801-B16 (START-grant).

## Author Contribution

JB and SN designed the research, SC performed bioinformatic and statistic analysis, JB and SC interpreted data; JB, SC and SN wrote the manuscript; All authors read and approved the final manuscript.

