## Supplementary Figures for "Dynamic cell wall modifications in brassicas during clubroot disease"

<u>**ELECTRONIC SUPPLEMENTARY MATERIAL**</u>

**Supplementary Notes**

Note S1: Data analysis p. 4
Note S2: Results of data set analysis p. 4

**Supplementary Tables**

**1. Biosynthesis of cell wall components**

Table S1: Regulation of gene expression coding for different cellulose synthase (CesA) genes p. 5

Table S2: Regulation of gene expression coding for different cellulose synthase –like (Csl) genes p. 6

Table S3: Regulation of gene expression coding for genes involved in pectin synthesis p. 7

Table S4: Regulation of gene expression coding for genes involved in lignin synthesis p. 8

Table S5: Regulation of gene expression coding for genes involved in callose synthesis and deposition p. 11

Table S6: Regulation of gene expression involved in sucrose metabolism p. 13

**2. Degradation and modification of cell wall components**

Table S1: Regulation of gene expression coding for pectin degrading and modifying enzymes like pectatlyases (PLs), pectin methylesterases (PMEs) and polygalacturonases (PGs) p. 14

Table S2: Regulation of gene expression coding for cellulose degrading and modifying enzymes p. 16

Table S3: Regulation of gene expression coding for enzyme inhibitors for pectin methylesterases (PMEs) and polygalacturonases (PGs) p. 19

**3. Cell wall loosening, elongation and response**

Table S4: Regulation of gene expression coding for xyloglucan endotransglucosylasen/hydrolasen p. 21

Table S5: Regulation of gene expression coding for expansins p. 22

Table S6: Regulation of wall associated kinases (WAKs) and wall associated kinase-like genes (WAKLs) p. 24

Table S7: Regulation of gene expression of Shaven3 (SHV3) and Shaven3 -like1 (SVL1) p. 25

Table S8: Regulation of gene expression of thioglucosidase (TGG) p. 26

Table S15: SI Data analysis – see excel file Suppl. Table S15 Additional file

### Note S1: Data analysis

Cell wall related genes were extracted from an existing *Brassica oleracea* var. *gongylodes* RNA-seq dataset (Ciaghi *et al.*, 2019a) (Accession: European Nucleotide Archive (ENA; https://www.ebi.ac.uk/ena) under the project PRJEB26435). These RNA-seq data originated from white spindle galls and brownish spindle galls and an uninfected control. Relevant pathways related to cell wall biosynthesis, rigidity, modulation, elongation, growth etc. were analyzed using all available information. Plant genes of interest (i.e. genes related with cellulose, hemicellulose, pectin, and lignin metabolism and cell expansion) were identified in this data using automatic gene annotations provided by MapMan (Thimm *et al.*, 2004), KAAS (KEGG Automatic Annotation Server; Moriya *et al.*, 2007), and eggNOG mapper (Huerta-Cepas *et al.*, 2016).

Transcripts fulfilling these selection criteria were curated manually. Only significantly differentially expressed genes (DEGs, adjusted p-value < 0.05) were analysed but some non-significant DEGs were included where it was useful for the biological interpretation and discussion of the data (noted in the text). This approach was chosen, because in a complex system like clubroots that contain a multitude of physiologically and developmentally different cells also smaller, not statistically significant changes might still be relevant for a biological discussion of the plant pathogen interaction and for the formulation of a biological hypothesis. Such instances were noted in the text, and many of the described pathways and pattern are found in most available datasets, yet functional validation is still missing and to provide this was beyond the scope of this review.

$Log_2$-fold change values of significant DEGs (infected vs. control) and their corresponding FPKM (fragments per kilobase per million reads) values were used for this study, without any further threshold values. MapMan, KAAS, eggNOG and TAIR Blast annotations were used to group genes of interest. Putative isoforms were validated using alignments of the amino acid and protein sequences, when contradictive expression values of DEGs were found. Isoforms were weighted based on their length, on sequence similarity and based on the general expression trend of all isoforms. Additionally, results published in studies on clubroot disease were integrated to provide a comprehensive overview on cell wall related changes in clubroot infected plants.

### Note S2: Results of data set analysis

Table S1 - S14: The following predicted **genes, functions** and **locations** are based on the information provided at The Arabidopsis Information Resource (TAIR, version 10) for the *Arabidopsis thaliana* homolog. All data refer to the RNA-seq dataset (Ciaghi *et al.*, 2019a) (Accession: European Nucleotide Archive (ENA; https://www.ebi.ac.uk/ena) under the project PRJEB26435. $Log_2$-fold change values of significant DEGs (infected vs. control)

### 1. Biosynthesis of cell wall components

#### Cellulose synthase GT2

Table S1: Regulation of gene expression coding for different cellulose synthase (CesA) genes in clubroot-infected roots compared to the control.

| Gene | Gene expression | Location | Function |
|---|---|---|---|
| CesA1 (at4g32410) | ↓ | Golgi apparatus, plasma membrane | cellulose synthase activity, transferase activity, transferring glycosyl groups |
| CesA2 (at4g39350) | ↓ | Plasmamembrane | |
| CesA3 (at5g05170) | ↓ | Golgi apparatus, plasma membrane | |
| CesA4 (at5g44030) | ↓ | Plasmamembrane | |
| CesA5 (at5g09870) | ↓ | membrane | |
| CesA6 (at5g64740) | ↓ | Golgi apparatus, plasma membrane, cortical microtubule | |

**Hemicellulose synthesis**

Table S2: Regulation of gene expression coding for different cellulose synthase –like (Csl) genes in clubroot-infected roots compared to the control.

| **Gene** | **Gene expression** | **Location** | **Function** |
|---|---|---|---|
| **Cellulose synthase-like** | | | |
| CSLA2 (at5g22740) | ↑ | | encodes a beta-mannan synthase |
| CSLC4 (at3g28180) | ↓ | plasma membrane | Might be involved in xyloglucan synthesis |
| CSLC5 (at4g31590) | ↑ | | |
| CSLC6 (at3g07330) | ↓ | Golgi apparatus | |
| CSLD2 (at5g16910) | ↑ | Golgi apparatus, plasma membrane, integral to Golgi membrane | Might be involved in mannansynthesis |
| CSLD3 (at3g03050) | ↓ | Golgi apparatus, endoplasmic reticulum, plasma membrane | Might be involved in mannansynthesis mutations initiate root hairs that rupture at their tip soon after initiation. is required for the synthesis of a noncellulosic wall polysaccharide |
| CSLD5 | ↑ | | Xylan, mannan synthesis |
| CSLE1 | ↑ | endoplasmic reticulum, plasma membrane | cellulose synthase activity, transferase activity, transferring glycosyl groups |

#### Pectin biosynthesis

Table S3: Regulation of gene expression coding for genes involved in pectin synthesis in clubroot-infected roots compared to the control.

| Gene | Gene expression | Location | Predicted Function |
|---|---|---|---|
| RHM2 (at1g53500) | ↓ | Chloroplast, cytosol | Encodes UDP-L-rhamnose synthase, required for rhamnoglacturonan1 synthesis |
| RHM1 (at1g78570) | ↓ | Chloroplast, cytosol, plasmodesma | Encodes UDP-L-Rhamnose synthase |
| QUA1 (at3g25140) | ↓ | Golgi apparatus, Golgi membrane, endosome, integral component of membrane, mitochondrion, trans-Golgi network | encodes a glycosyltransferase, involved in homogalacturonan biosynthesis |
| UXS1 (at3g53520) | ↑ | Golgi apparatus, Golgi cisterna membrane, cytoplasm, endosome, integral component of membrane, plasma membrane, trans-Golgi network | encodes an isoform of UDP-glucuronic acid decarboxylase, This enzyme produces UDP-xylose, which is a substrate for many cell wall carbohydrates including hemicellulose and pectin. UDP-xylose is also known to feedback regulate several cell wall biosynthetic enzymes |
| UXS2 (at3g62830) | ↓ | Golgi apparatus, Golgi cisterna membrane, cytoplasm, endosome, integral component of membrane, plasma membrane, trans-Golgi network | |
| UXS3 (at5g59290) | ↑ | cytoplasm, cytosol, plasma membrane | |
| UXS5 (at3g46440) | ↑ | cytoplasm, cytosol, plasma membrane | |

### Lignin biosynthesis

Table S4: Regulation of gene expression coding for genes involved in lignin synthesis in clubroot-infected roots compared to the control.

| Gene | Gene expression | Location | Function |
|---|---|---|---|
| **Lignin biosynthesis** | | | |
| OMT1 (at5g54160) | ↓ | cytosol, nucleus, plasma membrane, cytoplasm | myricetin 3-O-methyltransferase activity, quercetin 3-O-methyltransferase activity, caffeate O-methyltransferase activity |
| CCR1 (at1g76470) | ↑ | endomembrane system | coenzyme binding, binding, cinnamoyl-CoA reductase activity, catalytic activity |
| CCR2 (at1g80820) | ↓ | | |
| F5H, FAH1 (at4g36220) | ↓ | ER | ferulate 5-hydroxylase activity, monooxygenase activity |
| RPK2 (at3g02130) | ↑ | plasma membrane | protein serine/threonine kinase activity, kinase activity, ATP binding |
| BGLU46 (at1g61820) | ↓ | | cation binding, hydrolase activity, hydrolyzing O-glycosyl compounds, catalytic activity |
| MAT4, SAMS3, **MTO3** (at3g17390) | ↓ | nucleolus, cell wall, plasma membrane, membrane | methionine adenosyltransferase activity |
| LAC1 (at1g18140) | ↓ | endomembrane system, apoplast | |
| LAC3 (at2g30210) | ↓ | endomembrane system, apoplast | laccase activity |
| LAC6 (at2g46570) | ↑ | endomembrane system, apoplast | |
| LAC7 (at3g09220) | ↓ | endomembrane system, apoplast | |
| LAC11 (at5g03260) | ↓ | | |
| LAC13 (at5g07130) | ↑ | | |

| | | | |
|---|---|---|---|
| LAC17 (at5g60020) | ↓ | | |
| PA2 (at5g06720) | ↓ ↑ | Cell wall | peroxidase activity, heme binding |
| PRX34, PRX CB (at3g49120) | ↑ | apoplast, cell wall, plasma membrane, vacuole, plant-type cell wall | peroxidase activity, Involved in cell elongation |
| PRX37 (at4g08770) | ↑ | vacuole | response to oxidative stress, oxidation reduction |
| PRX72 (at5g66390) | ↓ | Endomembrane system | peroxidase activity, heme binding |
| at5g06730 | ↑ | Plasma membrane | |
| at5g06730 | ↑ | Plasma membrane | |
| at2g38390 | ↓ | | |
| at5g19890 | ↓ | nucleus, cytoplasm | |
| at2g38380 | ↓ | Cell wall | |
| at2g18150 | ↓ | Cell wall | |
| at3g32980 | ↓ | cell wall, vacuole, membrane | response to oxidative stress, oxidation reduction |
| **PAL** | | | |
| PAL1 (at2g37040) | ↓ | cytoplasm | phenylalanine ammonia-lyase activity |
| PAL2 (at3g53260) | ↓ | cytoplasm | |
| PAL4 (at3g10340) | ↓ | cytoplasm | |
| **C4H** | | | |
| (at2g30490) | ↓ | endoplasmic reticulum, plasma membrane, membrane, plant-type cell wall | trans-cinnamate 4-monooxygenase activity |
| **4CL** | | | 4-coumarate-CoA ligase activity |
| (at1g20480) | ↓ | | |
| (at1g20500) | ↑ | | |
| OPCL1 | ↑ | peroxisome | |

| | | | |
|---|---|---|---|
| (at1g20510) | | | |
| 4CL1 (at1g51680) | ↓ | | |
| 4CL5 (at3g21230) | ↓ | | |
| 4CL2 (at3g21240) | ↓ | | |
| C3H (at2g40890) | ↓ | mitochondrion, endoplasmic reticulum, plasma membrane, microsome | monooxygenase activity, p-coumarate 3-hydroxylase activity |

### Callose biosynthesis, -binding and -deposition

Table S5: Regulation of gene expression coding for genes involved in callose synthesis –binding and deposition in clubroot-infected roots compared to the control.

<table>
<tr><th>Gene:</th><th>Gene expression</th><th>Location</th><th>Function</th></tr>
<tr><td>Callose biosynthesis</td><td></td><td></td><td></td></tr>
<tr><td>CalS1<br>at1g05570</td><td>↑</td><td></td><td>Encodes a callose synthase 1 catalytic subunit</td></tr>
<tr><td>GSL1<br>at4g04970</td><td>↑</td><td rowspan="6">1,3-beta-glucan synthase complex, plasma membrane</td><td rowspan="6">transferase activity, transferring glycosyl groups, 1,3-beta-glucan synthase activity</td></tr>
<tr><td>GSL03<br>at2g31960</td><td>↓</td></tr>
<tr><td>GSL07<br>at1g06490</td><td>↓</td></tr>
<tr><td>GSL8<br>at2g36850</td><td>↓</td></tr>
<tr><td>GSL10<br>at3g07160</td><td>↓</td></tr>
<tr><td>GSL12<br>at5g13000</td><td>↓</td></tr>
<tr><td>UGT1<br>at1g05560</td><td>↓</td><td></td><td></td></tr>
<tr><td>Callose binding</td><td></td><td></td><td></td></tr>
<tr><td>PDCB1<br>at5g61130</td><td>↑</td><td rowspan="4">Cell wall, plasmodesma, anchored to plasma membrane, plasma membrane, anchored to membrane</td><td>callose binding, polysaccharide binding, overexpression leads to accumulation of callose</td></tr>
<tr><td>PDCB3<br>at1g18650</td><td>↑</td><td rowspan="3">callose binding, polysaccharide binding</td></tr>
<tr><td>PDCB4<br>at1g69295</td><td>↓</td></tr>
<tr><td>PDCB4<br>at3g58100</td><td>↑</td></tr>
<tr><td>Callose deposition</td><td></td><td></td><td></td></tr>
</table>

| | | | |
|---|---|---|---|
| BBD1<br>at1g75380 | ↑ | | |
| UGP2<br>at5g17310 | ↑ | plasma membrane | UTP:glucose-1-phosphate uridylyltransferase activity, nucleotidyltransferase activity |
| PEN2<br>at2g44490 | ↑ | peroxisome, membrane, chloroplast envelope; | thioglucosidase activity, hydrolase activity, hydrolyzing O-glycosyl compounds; involved in: defense response to fungus, incompatible interaction, glucosinolate metabolic process, defense response to bacterium, indole glucosinolate catabolic process, defense response by callose deposition in cell wall |
| TSA1/2<br>at3g54640 | ↑ | chloroplast | tryptophan synthase activity; involved in: tryptophan biosynthetic process, defense response to bacterium, defense response by callose deposition in cell wall |
| AGD2<br>at4g33680 | ↑ | chloroplast, chloroplast stroma | transaminase activity, copper ion binding, L,L-diaminopimelate aminotransferase activity |
| VTC2<br>at4g26850 | ↑ | | |

### Sugar metabolism

Table S6: Regulation of gene expression involved in sucrose metabolism in clubroot-infected roots compared to the control.

| Gene | Gene expression | Location | Function |
|---|---|---|---|
| SUS1 (at5g20830) | ↑ | | Sucrose synthase activity |
| SUS5 (at5g37180) | ↑ | | Sucrose synthase activity |
| SPS4F (at4g10120) | ↑ | plasmamembrane | Sucrose synthesis |
| SPS2F (at5g11110) | ↑ | plasmamembrane | Sucrose synthesis |
| SPP1 (at1g51420) | ↑ | plasmamembrane | Sucrose synthesis |

### 2. Degradation and modification of cell wall components

#### Pectatlyases, pectin methylesterases, polygalacturonases

Table S7: Regulation of gene expression coding for pectin degrading and modifying enzymes like pectatlyases (PLs), pectin methylesterases (PMEs) and polygalacturonases (PGs) in clubroot-infected roots compared to the control.

| **Gene** | **Enzyme** | **Gene expression** | **Location** | **Function** |
|---|---|---|---|---|
| PMR6 (at3g54920) | **PL** | ↑ | anchored component of membrane, anchored component of plasma membrane, plasma membrane | |
| PLL19 (at4g24780) | **PL** | ↑ | | Pectin lyase-like superfamily protein |
| at1g67750 | **PL** | ↑ | endomembrane system | pectate lyase activity |
| at1g04680 | **PL** | ↑ | membrane | Pectin lyase-like superfamily protein, pectate lyase activity |
| at4g13710 | **PL** | ↑ | endomembrane system | Pectin lyase-like superfamily protein; lyase activity, pectate lyase activity |
| at3g09540 | **PL** | ↓ | | Pectin lyase-like superfamily protein; lyase activity, pectate lyase activity |
| at3g24670 | **PL** | ↓ | Endomembrane system | Pectin lyase-like superfamily protein; pectate lyase activity |
| at3g27400 | **PL** | ↓ | Endomembrane system | Pectin lyase-like superfamily protein; pectate lyase activity |
| at5g04310 | **PL** | ↑ | anchored component of membrane | Pectin lyase-like superfamily protein |
| at5g63180 | **PL** | ↑ | | Pectin lyase-like superfamily protein |
| at3g53190 | **PL** | ↑ | anchored component of membrane, transcellular region | Pectin lyase-like superfamily protein |
| at1g10640 | **PG** | ↑ | Extracellular region | Pectin lyase-like superfamily protein; polygalacturonase activity |
| at1g19170 | **PG** | ↓ | chloroplast | |
| at1g23460 | **PG** | ↓ | Endomembrane system | |
| at1g80170 | **PG** | ↑ | Cell wall | |
| at2g43890 | **PG** | ↑ | endomembrane system | |
| at4g23820 | **PG** | ↑ | Endomembrane system | |
| at5g41870 | **PG** | ↑ | Endomembrane system | |

| | | | | |
|---|---|---|---|---|
| at1g02460 | **PG** | ↓ | | Pectin lyase-like superfamily protein; polygalacturonase activity |
| at3g07970 | **PG** | ↓ | endomembrane system | May be required for cell type-specific pectin degradation. polygalacturonase activity |
| at3g16850 | **GH28** | ↑ | Plant cell wall, vacuole | |
| at3g61490 | **GH28** | ↑ | Endomembrane system, chloroplast | Pectin lyase-like superfamily protein; polygalacturonase activity |
| at5g49215 | **GH28** | ↑ | Endomembrane system | |
| **PMEs** | | | | |
| PMEPCRA (at1g11580) | | ↑ | cell wall, plasma membrane, plant-type cell wall | enzyme inhibitor activity, pectinesterase activity |
| SKS6 (at1g41830) | | ↑ | apoplast, cell wall, membrane, plant-type cell wall | pectinesterase activity |
| PAE12 (at3g05910) | | ↓ | | carboxylesterase activity |
| PAE5 (at3g09410) | | ↓ | | carboxylesterase activity |
| PME3 (at3g14310) | | ↑ | cell wall, apoplast, plasma membrane, cytoplasm | pectinesterase activity |
| PME31 (at3g29090) | | ↑ | cell wall, plant-type cell wall | pectinesterase activity |
| PAE7 (at4g19410) | | ↑ | Cell wall | actin binding, carboxylesterase activity |
| at5g19730 | | ↑ | Cell wall, cytoplasm | pectinesterase activity |
| PAE9 (at5g23870) | | ↑ | Cell wall | carboxylesterase activity |
| PAE10 (at5g26670) | | ↑ | | Pectin acetylesterase activity |
| PAE11 (at5g45280) | | ↑ | Cell wall | carboxylesterase activity |
| PME5 (at5g47500) | | ↑ | endomembrane system, cell wall, plant-type cell wall | pectinesterase activity |

### Cellulasen

Table S8: Regulation of gene expression coding for cellulose degrading and modifying enzymes in clubroot-infected roots compared to the control.

<table>
<tr><th>Gene</th><th>Gene expression</th><th>Location</th><th>Function</th></tr>
<tr><td>GH9B1 (at1g70710)</td><td>↑</td><td>chloroplast</td><td>cellulase activity, hydrolase activity, hydrolyzing O-glycosyl compounds, INVOLVED IN: response to cyclopentenone, cell wall modification involved in multidimensional cell growth</td></tr>
<tr><td>GH9B6 (at1g23210)</td><td>↓</td><td>endomembrane system</td><td rowspan="5">hydrolase activity, hydrolyzing O-glycosyl compounds, catalytic activity</td></tr>
<tr><td>GH9B7 (at1g75680)</td><td>↑</td><td>Plasma membrane</td></tr>
<tr><td>GH9B8 (at2g32990)</td><td>↑</td><td>endomembrane system</td></tr>
<tr><td>GH9B12 (at2g44570)</td><td>↑</td><td>endomembrane system</td></tr>
<tr><td>GH9B13 (at4g02290)</td><td>↓</td><td>endomembrane system</td></tr>
<tr><td>GH9A1 (at5g49720)</td><td>↓</td><td>Golgi apparatus, plasma membrane, cell plate, early endosome</td><td>cellulase activity, hydrolase activity, hydrolyzing O-glycosyl compounds;</td></tr>
<tr><td>GH9C3 (at4g11050)</td><td>↓</td><td>endomembrane system, extracellular region</td><td>carbohydrate binding, hydrolase activity, hydrolyzing O-glycosyl compounds, catalytic activity</td></tr>
<tr><td>BGLU8 (at3g62750)</td><td>↓</td><td>vacuole</td><td>Cation binding, hydrolase activity, hydrolyzing O-glycosyl compounds</td></tr>
<tr><td>BGLU9 (at4g27820)</td><td>↓</td><td>peroxisome</td><td>Cation binding, hydrolase activity, hydrolyzing O-glycosyl compounds</td></tr>
<tr><td>BGLU10 (at4g27830)</td><td>↓</td><td>vacuole</td><td>cation binding, hydrolase activity, hydrolyzing O-glycosyl compounds, catalytic activity</td></tr>
<tr><td>BG3 (at3g57240)</td><td>↑</td><td>Cell wall, apoplast</td><td>cellulase activity, hydrolase activity, hydrolyzing O-glycosyl compounds</td></tr>
<tr><td>BGL2 (at3g57260)</td><td>↑</td><td>apoplast, cell wall, vacuole</td><td>glucan 1,3-beta-glucosidase activity, protein binding, cellulase activity, hydrolase activity, hydrolyzing O-glycosyl compounds/</td></tr>
</table>

| | | | |
|---|---|---|---|
| | | | systemic acquired resistance, response to cold |
| GH family17 (at2g16230) | ↑ | endomembrane system | |
| GH family17 (at2g05790) | ↑ | endomembrane system | |
| GH family17 (at2g27500) | ↓ | anchored to plasma membrane, nucleus, cytoplasm | |
| GH family17 (at1g11820) | ↑ | endomembrane system | |
| GH family17 (at3g13560) | ↑ | anchored to plasma membrane, plasma membrane, anchored to membrane | |
| GH family17 (at5g58090) | ↑ | anchored to plasma membrane, plasma membrane, anchored to membrane | |
| GH family17 (at2g26600) | ↓ | anchored to membrane | |
| GH family17 (at3g55430) | ↓ | Cell wall | cation binding, hydrolase activity, hydrolyzing O-glycosyl compounds, catalytic activity |
| GH family17 (at2g16230) | ↑ | Endomembrane system | |
| GH family 17 (at5g58480) | ↑ | anchored to plasma membrane, plasma membrane, anchored to membrane | |
| GH family 17 (at3g55430) | ↑ | Cell wall | |
| GH family 17 (at4g31140) | ↓ | anchored to plasma membrane, plasma membrane, anchored to membrane | |
| GH family 17 (at4g29360) | ↑ | anchored to plasma membrane, plasma membrane, anchored to membrane | |
| GH family 17 (at4g17180) | ↑ | endomembrane system | |
| GH family 17 (at5g55180) | ↓ | endomembrane system | |
| at5g20950 | ↑ | cell wall, membrane, plant-type cell wall | Beta-glucosidase involved in xyloglucan metabolism |
| at3g47000 | ↑ | Chloroplast, cytoplasm | hydrolase activity, hydrolyzing O-glycosyl compounds |
| at3g47010 | ↑ | Chloroplast, cytoplasm | hydrolase activity, hydrolyzing O-glycosyl compounds |

| | | | |
|---|---|---|---|
| at3g47040 | ↓ | | hydrolase activity, hydrolyzing O-glycosyl compounds |
| at2g27500 | ↑ | anchored to plasma membrane, nucleus, cytoplasm | cation binding, hydrolase activity, hydrolyzing O-glycosyl compounds, catalytic activity |
| at1g32860 | ↓ | anchored to plasma membrane, nucleus, cytoplasm | |
| at1g66250 | ↑ | anchored to plasma membrane, plasma membrane, anchored to membrane | |
| at4g18340 | ↓ | endomembrane system | |
| at4g31140 | ↓ | anchored to plasma membrane, plasma membrane, anchored to membrane | |
| **callose degradation** | | | |
| at5g42100 | ↑ | | hydrolase activity, hydrolyzing O-glycosyl compounds, glucan endo-1,3-beta-D-glucosidase activity |

### Inhibitors

Table S9: Regulation of gene expression coding for enzyme inhibitors for pectin methylesterases (PMEs) and polygalacturonases (PGs) in clubroot-infected roots compared to the control.

| **Gene** | **Gene expression** | **Location** | **Function** |
|---|---|---|---|
| PGIP1 (at5g06860) | ↓ ↑ | Golgi apparatus, cell wall, extracellular region, membrane, plant-type cell wall, plasmodesma | Encodes a polygalacturonase inhibiting protein involved in defense response. PGIPs inhibit the function of cell wall pectin degrading enzymes such as those produced by fungal pathogens. PGIP1 is induced by fungal infection. |
| PGIP2 (at5g06870) | ↓ | cell wall, extracellular region, membrane, plant-type cell; wall, plasmodesma, not in cytosol | Encodes a polygalacturonase inhibiting protein involved in plant defense response. PGIPs inhibit the activity of pectin degrading enzymes such as those produced by fungal pathogens. PGIP2 is induced by fungal infection and methyl jasmonate.; polygalacturonase inhibiting protein 2 (PGIP2) |
| **PMEI** | | | |
| RHS12 root-hair-specific (at3g10710) | ↓ | | enzyme inhibitor activity, pectinesterase activity |
| PMEI6 (at2g47670) | ↓ | endomembrane system | enzyme inhibitor activity, pectinesterase inhibitor activity, pectinesterase activity |
| PMEI7 (at4g2526) | ↑ | endomembrane system | |
| PMEI11 (at3g47380) | ↓ | endomembrane system | |
| PMEI13 (at5g62360) | ↓ | | |
| PME17 (at2g45220) | ↑ | membrane, plant-type cell wall | enzyme inhibitor activity, pectinesterase activity |
| PME34 (at3g49220) | ↑ | Cell wall | enzyme inhibitor activity, pectinesterase activity |
| PME35/61 (at3g59010) | ↓ | | enzyme inhibitor activity, pectinesterase activity<br>PME35-mediated demethylesterification of the primary cell wall regulates the mechanical strength of the supporting tissue. |

| | | | |
|---|---|---|---|
| C/VIF2 (at5g64620) | ↑ | endomembrane system, chloroplast, vacuole | enzyme inhibitor activity, pectinesterase inhibitor activity, pectinesterase activity |
| C/VIF1 (at1g47960) | ↑ | Cell wall | enzyme inhibitor activity, pectinesterase inhibitor activity, pectinesterase activity, Inhibits Invertase/fructosidase |
| at5g04970 | ↓ | | enzyme inhibitor activity, pectinesterase activity |
| at2g47550 | ↓ | endomembrane system, cell wall, plant-type cell wall | |
| at3g10720 | ↑ | endomembrane system, cell wall, plant-type cell wall | |
| at3g17130 | ↑ | endomembrane system | enzyme inhibitor activity, pectinesterase inhibitor activity, pectinesterase activity |
| at3g43270 | ↓ | Cell wall | enzyme inhibitor activity, pectinesterase activity |
| at3g47670 | ↑ | Golgi apparatus, chloroplast | enzyme inhibitor activity, pectinesterase inhibitor activity, pectinesterase activity |
| at5g09760 | ↑ | cell wall, chloroplast | enzyme inhibitor activity, pectinesterase activity |
| at5g38610 | ↑ | endomembrane system | enzyme inhibitor activity, pectinesterase inhibitor activity, pectinesterase activity |
| at5g62350 | ↓ | endomembrane system | enzyme inhibitor activity, pectinesterase inhibitor activity, pectinesterase activity |
| at1g02810 | ↑ | endomembrane system, cell wall | enzyme inhibitor activity, pectinesterase activity |
| at1g14890 | ↓ | mitochondrion | enzyme inhibitor activity, pectinesterase inhibitor activity, pectinesterase activity |

### 3. Cell wall loosening, elongation and response

#### XTH

Table S10: Regulation of gene expression coding for xyloglucan endotransglucosylasen/hydrolasen in clubroot-infected roots compared to the control.

<table>
<tr><th>Gene</th><th>Gene expression</th><th>Location</th><th>Function</th></tr>
<tr><td>XTH 4 (at2g06850)</td><td>↑</td><td></td><td>hydrolase activity, acting on glycosyl bonds, xyloglucan:xyloglucosyl transferase activity, Involved in response to auxin stimulus, response to mechanical stimulus, response to low light intensity stimulus, unidimensional cell growth</td></tr>
<tr><td>XTH 9 (at4g03210)</td><td>↑</td><td>apoplast, cell wall</td><td>hydrolase activity, acting on glycosyl bonds, xyloglucan:xyloglucosyl transferase activity</td></tr>
<tr><td>XTH 16 (at3g23730)</td><td>↑</td><td>endomembrane system, cell wall, apoplast</td><td>hydrolase activity, acting on glycosyl bonds, hydrolase activity, hydrolyzing O-glycosyl compounds, xyloglucan:xyloglucosyl transferase activity</td></tr>
<tr><td>XTH 24 (at4g30270)</td><td>↑</td><td>cell wall, plasma membrane, cytoplasm</td><td>hydrolase activity, acting on glycosyl bonds, xyloglucan:xyloglucosyl transferase activity, Involved in: aging, response to gibberellin stimulus, gibberellic acid mediated signaling pathway, response to brassinosteroid stimulus, plant-type cell wall loosening</td></tr>
<tr><td>XTH 32 (at2g36870)</td><td>↑</td><td>endomembrane system</td><td>hydrolase activity, acting on glycosyl bonds, hydrolase activity, hydrolyzing O-glycosyl compounds</td></tr>
<tr><td>XTH 8 (at1g11545)</td><td>↓</td><td>endomembrane system, cell wall, apoplast</td><td>hydrolase activity, acting on glycosyl bonds, xyloglucan:xyloglucosyl transferase activity, hydrolase activity, hydrolyzing O-glycosyl compounds</td></tr>
<tr><td>XTH 10 (at2g14620)</td><td>↓</td><td>endomembrane system, apoplast, cell wall;</td><td>hydrolase activity, acting on glycosyl bonds, xyloglucan:xyloglucosyl transferase activity, hydrolase activity, hydrolyzing O-glycosyl compounds</td></tr>
<tr><td>XTH 21 (at2g18800)</td><td>↓</td><td>endomembrane system, apoplast, cell wall</td><td rowspan="3">hydrolase activity, acting on glycosyl bonds, xyloglucan:xyloglucosyl transferase activity, hydrolase activity, hydrolyzing O-glycosyl compounds</td></tr>
<tr><td>XET, XTH 18 (at4g30280)</td><td>↓</td><td>endomembrane system, apoplast, cell wall</td></tr>
<tr><td>XTR 8 (at3g44990)</td><td>↓</td><td>endomembrane system, cell wall, apoplast</td></tr>
</table>

**Expansins**

Table S11: Regulation of gene expression coding for expansins in clubroot-infected roots compared to the control.

| Gene | Gene expression | Location | Function |
|---|---|---|---|
| EXP1 (at1g69530) | ↑ | | |
| EXP3 (at2g37640) | ↓ | endomembrane system, extracellular region | |
| EXP4 (at2g39700) | ↓ | | |
| EXP6 (at2g28950) | ↑ | | |
| EXP9 (at5g02260) | ↓ | chloroplast | response to cyclopentenone, plant-type cell wall organization, unidimensional cell growth, plant-type cell wall loosening |
| EXP13 (at3g03220) | ↑ | extracellular region | response to cyclopentenone, plant-type cell wall organization, unidimensional cell growth, plant-type cell wall loosening |
| EXP15 (at2g03090) | ↑ | endomembrane system, extracellular region | plant-type cell wall modification involved in multidimensional cell growth, syncytium formation, unidimensional cell growth, plant-type cell wall loosening |
| EXP17 (at4g01630) | ↓ | endomembrane system, extracellular region | plant-type cell wall modification involved in multidimensional cell growth, unidimensional cell growth, plant-type cell wall loosening |
| EXP20 (at4g38210) | ↑ | endomembrane system, extracellular region | Ivolved in plant-type cell wall modification involved in multidimensional cell growth, response to karrikin, syncytium formation, unidimensional cell growth, plant-type cell wall loosening |
| EXLA1 (at3g45970) | ↓ | Cell wall | unidimensional cell growth, plant-type cell wall loosening |
| EXLA2 (at4g38400) | ↓ | endomembrane system, extracellular region | response to cyclopentenone, plant-type cell wall organization, unidimensional cell growth, plant-type cell wall loosening |
| EXLB1 (at4g17030) | ↑ | endomembrane system, extracellular region | sexual reproduction, unidimensional cell growth, plant-type cell wall loosening |

| | | | |
|---|---|---|---|
| EXPB2 (at1g65680) | ↑ | endomembrane system, extracellular region | response to cyclopentenone, plant-type cell wall organization, unidimensional cell growth, plant-type cell wall loosening |
| EXPB3 (at4g28250) | ↑ | endomembrane system, extracellular region | response to cyclopentenone, plant-type cell wall organization, unidimensional cell growth, plant-type cell wall loosening |

### WAKs/WAKLs

Table S12: Regulation of wall associated kinases (WAKs) and wall associated kinase-like genes (WAKLs) in clubroot-infected roots compared to the control.

| Gene | Gene expression | Location | Funktion |
|---|---|---|---|
| WAK (at1g17910) | ↑ | endomembrane system, integral to membrane | kinase activity |
| WAK (at1g16260) | ↑ | endomembrane system, integral to membrane | kinase activity |
| WAK4 (at1g21210) | ↑ | plasma membrane | protein serine/threonine kinase activity, protein kinase activity, ATP binding, calcium ion binding |
| WAKL (at5g66790) | ↓ | endomembrane system | kinase activity |
| WAKL4 (at1g16150) | ↓ | Cell wall, plasma membrane, cell surface | Encodes a WAK-like receptor-like kinase with a cytoplasmic Ser/Thr protein kinase domain and an extracellular domain with EGF-like repeats |
| WAKL (at2g23450) | ↓ | Integral component of membrane, plasma membrane | kinase activity |

### Shaven3 (SHV3) and Shaven3 -like1 (SVL1)

Table S13: Regulation of gene expression of Shaven3 (SHV3) and Shaven3 -like1 (SVL1) in clubroot-infected roots compared to the control.

| Gene | Gene expression | Location | Funktion |
|---|---|---|---|
| GDPDL3, SHV3 (at4g26690) | ↑ | anchored to plasma membrane, plasma membrane, anchored to membrane, plant-type cell wall | glycerophosphodiester phosphodiesterase activity, kinase activity |
| GDPDL4, SVL1 (at5g55480) | ↑ | anchored to plasma membrane, plasma membrane, anchored to membrane | involved in cell wall cellulose accumulation and pectin linking phosphoric diester hydrolase activity, glycerophosphodiester phosphodiesterase activity |

### Thioglucosidase (TGG)

Table S14: Regulation of gene expression of thioglucosidase (TGG) in clubroot-infected roots compared to the control.

| Gene | Gene expression | Location | Funktion |
|---|---|---|---|
| TGG1 (at5g26000) | ↓ | | thioglucosidase activity, beta-glucosidase activity, hydrolase activity, hydrolyzing O-glycosyl compounds |
| TGG2 (at5g25980) | ↓ | cytosolic ribosome, apoplast, chloroplast, peroxisome, vacuole | thioglucosidase activity, hydrolase activity, hydrolyzing O-glycosyl compounds |
